## Supplementary material for "Regulatory T cells suppress the formation of potent KLRK1 and IL-7R expressing effector CD8 T cells by limiting IL-2": Figure Supplement

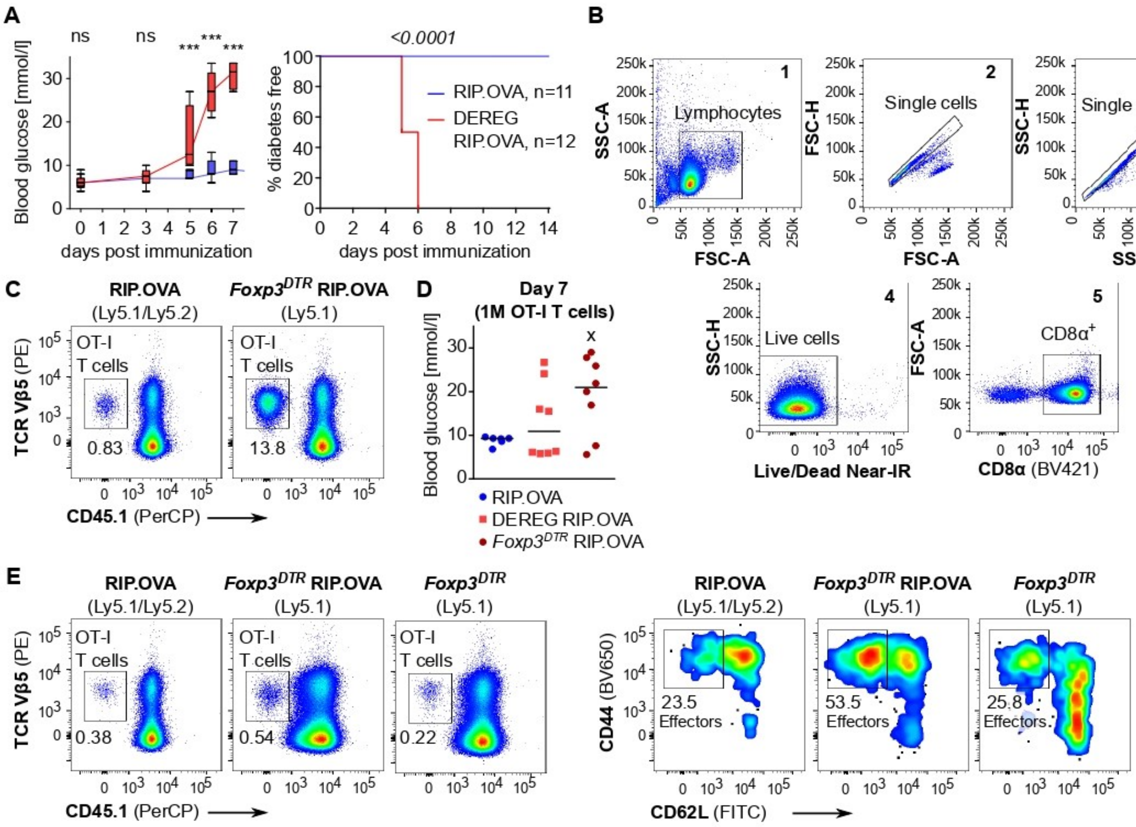


**Figure 1 – Figure supplement 1**

A. Diabetes was induced as shown in Figure 1A-C with 1×104 OT-I T cells in two independent experiments in RIP.OVA (n=11) and DEREG RIP.OVA mice (n=12). Left: glucose concentration in blood was recorded on days 0, 3, 5, 6, 7 post OVA + LPS injection. Statistical significance was calculated by two-tailed Mann-Whitney test. P-value ***<0.001. Median and range is shown. Right: glucose concentration in urine was monitored on a daily basis. Percentage of diabetes free mice in time is shown. Statistical significance was calculated by Log-rank (Mantel-Cox) test, p-value is shown in italics.

B. Gating strategy for the experiment shown in Figure 1F. CD4+ T cells and B cells were depleted via magnetic bead separation prior to the analysis.

C. Representative dot plots for the experiment shown in Figure 1F.

D. Blood glucose concentration (day 7) in the experiment shown in Figure 1G (106 transferred OT-I T cells). RIP.OVA n=6, DEREG RIP.OVA n=8, Foxp3DTR RIP.OVA n=8. One Foxp3DTR RIP.OVA mouse died before the measurement (shown as “x”). Median is shown.

E. Representative dot plots for the experiment shown in Figure 1H.

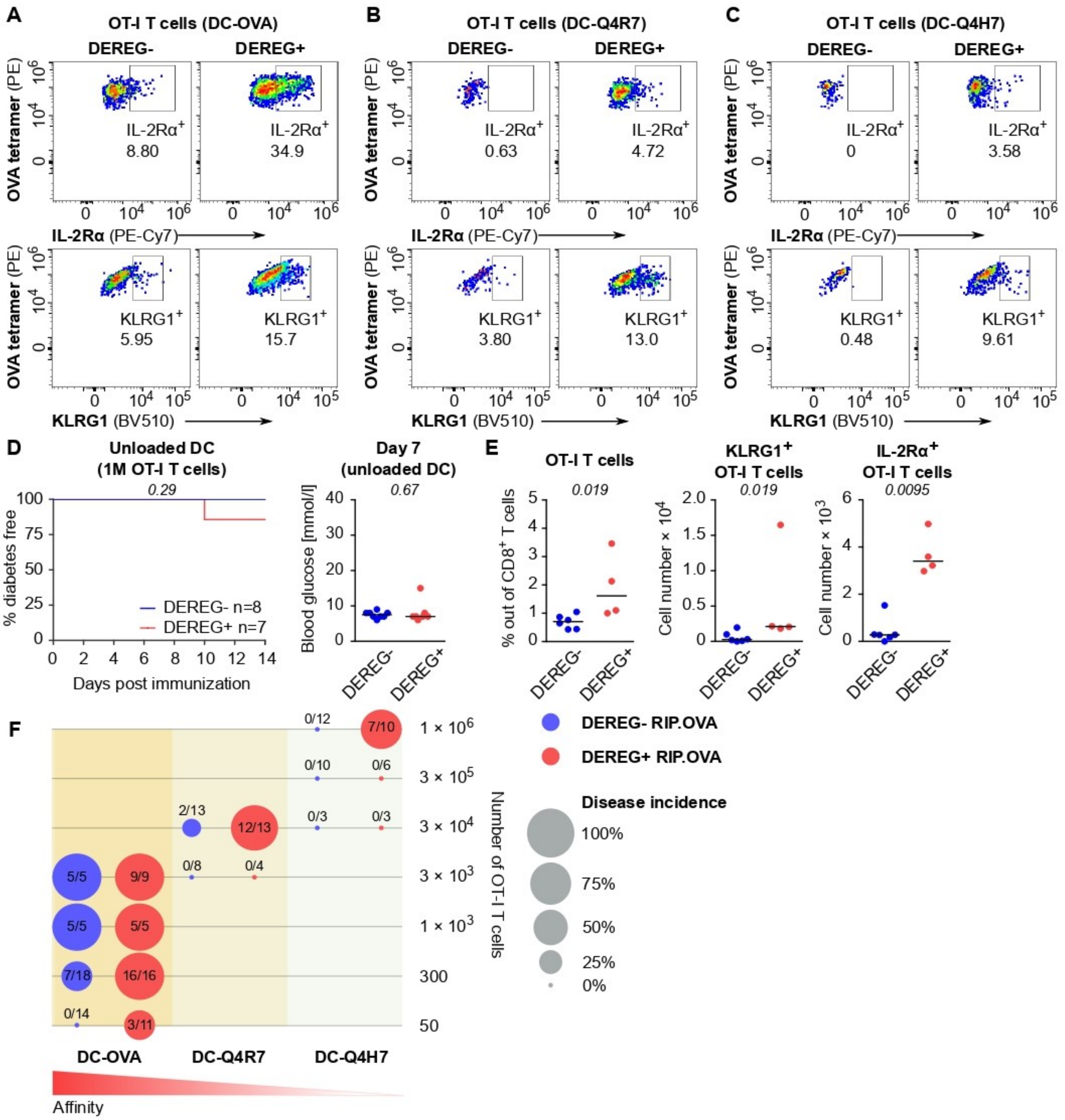


**Figure 2 – Figure supplement 1**

**A.** Representative dot plots for the experiment shown in Figure 2D. **B.** Representative dot plots for the experiment shown in Figure 2E. **C.** Representative dot plots for the experiment shown in Figure 2F. **D-E.** Treg-depleted DEREG$^+$ RIP.OVA mice and control DEREG$^-$ RIP.OVA mice received $10^6$ OT-I T cells. The next day, mice were immunized with unloaded DC. **D.** Urine glucose level was monitored on a daily basis. Left: Percentage of diabetes-free mice. Number of mice per group is indicated. Right: Blood glucose concentration on day 7 post- immunization. **E.** On day 6 post-immunization, spleens were collected and analyzed by flow cytometry. Percentage of OT-I T cells among CD8$^+$ T cells, count of KLRG1$^+$ OT-I T cells, and count of IL-2Rα$^+$ OT-I T cells are shown. DEREG$^-$ n=6, DEREG$^+$ n=4.

**F.** Bubble chart illustrating diabetes incidence in DEREG$^+$ RIP.OVA and DEREG$^-$ RIP.OVA mice immunized with DC loaded with indicated peptides vs. numbers of transferred OT-I T cells. Number of diabetic mice and total number of mice per group is indicated.

Statistical significance was calculated by Log-rank (Mantel-Cox) test (survival) or two-tailed Mann-Whitney test (glucose concentration and flow cytometry analysis), p-value is shown in italics. Median is shown.

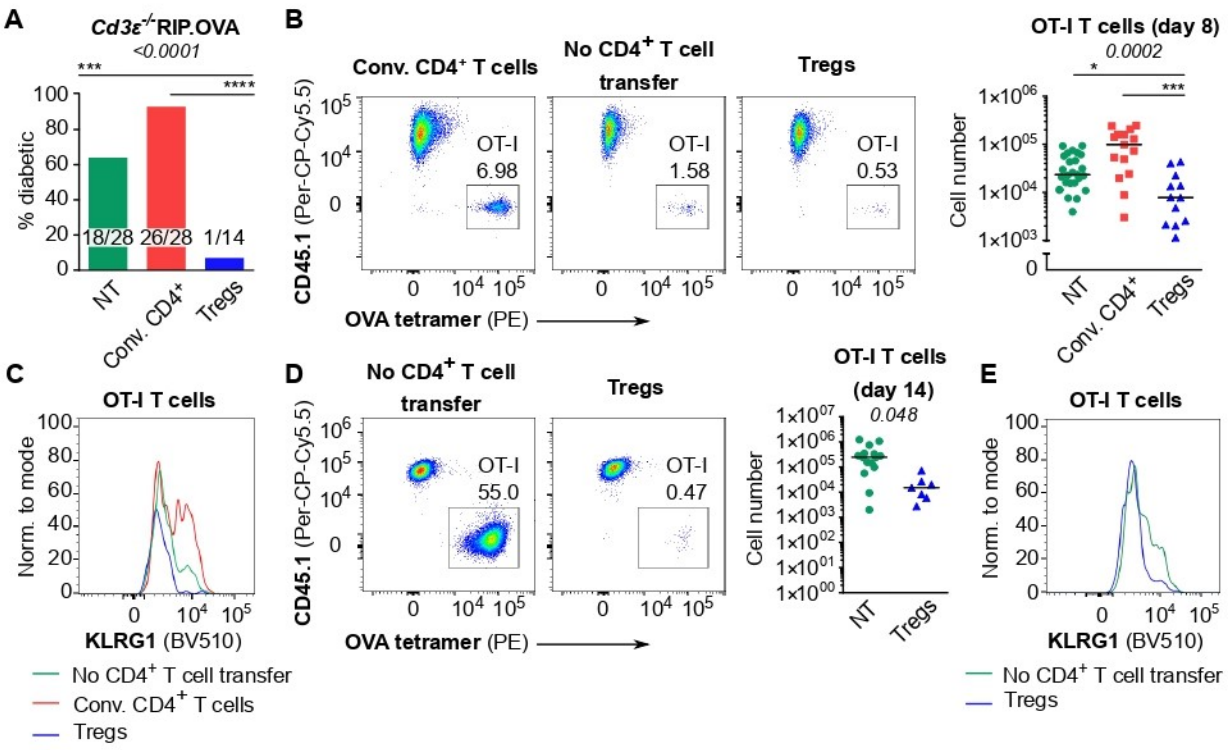


**Figure 3 – Figure supplement 1**

**A.** Percentage of diabetic mice in individual groups in the experiment shown in Figure 3A. Number of diabetic mice and total number of mice per group is indicated.

**B-C.** In the experimental setup shown in Figure 3A, spleens were collected on day 8 post-immunization and analyzed by flow cytometry. **B.** Left: Percentage of OT-I T cells among $CD8^+$ T cells. A representative experiment out of five in total. Right: Number of OT-I T cells is shown. NT n=25, Conv. $CD4^+$ n=15, Tregs n=12. **C.** KLRG1 expression on OT-I T cells on day 8, a representative histogram.

**D-E.** In the experimental setup shown in Figure 3A, spleens were collected on day 14 post-immunization and analyzed by flow cytometry. **D.** Left: Percentage of OT-I T cells among $CD8^+$ T cells. Representative plots are shown. Right: Number of OT-I T cells is shown. NT n=15, Tregs n=7. **E.** KLRG1 expression on OT-I T cells on day 14. A representative experiment out of 3 in total.

Statistical significance was calculated using Chi-square test (p-value is shown in italics) and Fisher's exact post-test (***<0.001, ****<0.0001) for comparison of two groups of the outcomes: diabetic mice vs. diabetes-free mice (**A**). Statistical significance for flow cytometry analysis was calculated by Kruskal-Wallis test (p-value is shown in italics) with Dunn's post-test (* <0.05, ***<0.001, ****<0.0001) (**B**), or two-tailed Mann-Whitney test (**D**). Median is shown.

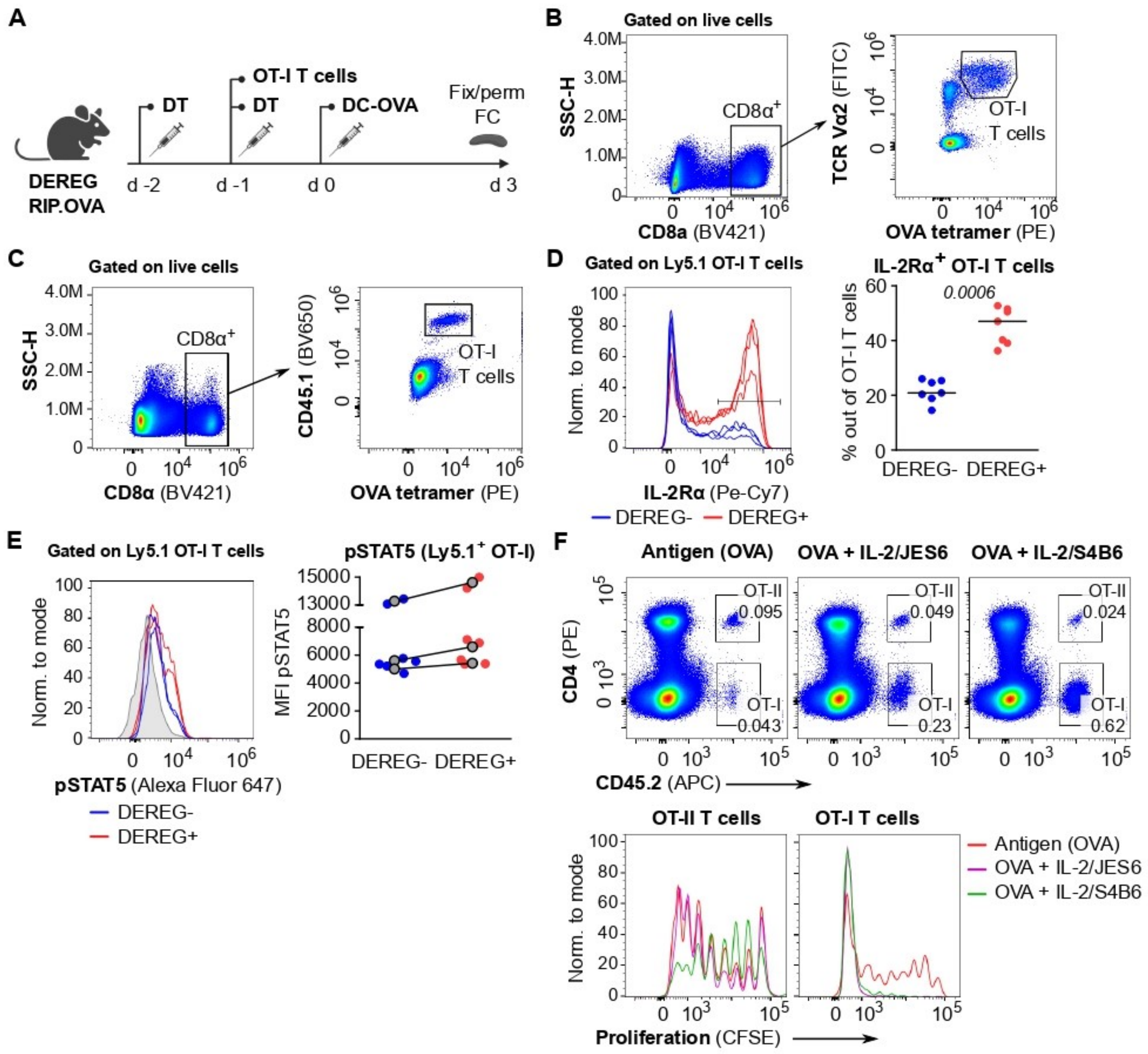


**Figure 4 – Figure supplement 1**

**A.** Scheme of the experiments described in Figure 4A-C and Figure 4 supplement 1B-E.
**B.** Gating strategy for the experiment shown in Figure 4A.
**C-E.** The experiment described in Figure 4A was modified by using congenic Ly5.1 OT-I T cells for the adoptive transfer. **C.** The gating strategy. **D.** IL-2Rα expression on OT-I Ly5.1 T cells. Left: a representative experiment out of 3 in total. Right: Percentage of IL-2Rα$^+$ cells among OT-I Ly5.1 T cells, n=7 mice per group. Statistical significance was calculated by two-tailed Mann-Whitney test, p-value is shown in italics. Median is shown. **E.** pSTAT5 expression in OT-I Ly5.1 T cells. Left: a representative experiment out of 3 in total. Right: Geometric mean fluorescence intensity (MFI) of anti-pSTAT5-Alexa Fluor 647 on Ly5.1 OT-I T cells. Mean of MFI values for each genotype per experiment are shown as a grey dots. Lines connect data from corresponding experiments. n=7 mice per group.

**F.** OT-I CD8$^+$ and OT-II CD4$^+$ T cells (both Ly5.2) were labeled with CFSE and adoptively transferred into Ly5.1 mice as a mixture ($0.75 \times 10^6$ CD8$^+$ OT-I + $1.5 \times 10^6$ CD4$^+$ OT-II T cells), followed by the administration of OVA protein with or without IL-2ic (IL-2/S4B6 or IL-2/JES6). On day 5 post-immunization, spleens were collected and analyzed by flow cytometry. Top: Representative staining showing CD45.2$^{+}$ OT-I (CD4$^{-}$) and OT-II (CD4$^{+}$) T cells among viable cells. Bottom: Proliferation is indicated by the CFSE dilution. Representative histograms for OT-I and OT-II T cells.

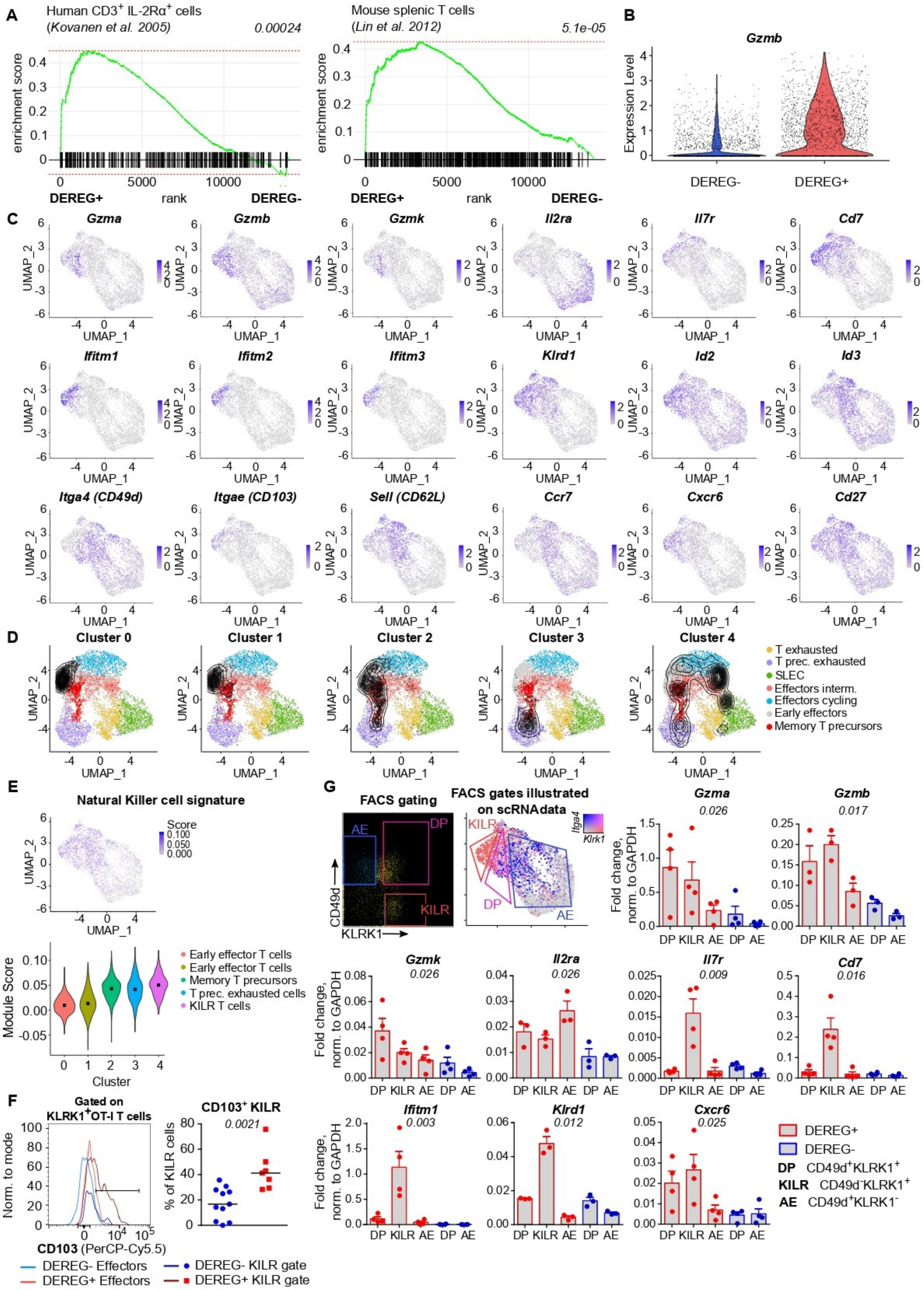
A
Human CD3+ IL-2Rα+ cells
(Kovanen et al. 2005)
0.00024
enrichment score
rank
DEREG+
DEREG-
Mouse splenic T cells
(Lin et al. 2012)
5.1e-05
B
Gzmb
Expression Level
DEREG-
DEREG+
C
Gzma
Gzmb
Gzmk
Il2ra
Il7r
Cd7
Ifitm1
Ifitm2
Ifitm3
Klrd1
Id2
Id3
Itga4 (CD49d)
Itgae (CD103)
Sell (CD62L)
Ccr7
Cxcr6
Cd27
UMAP_1
UMAP_2
D
Cluster 0
Cluster 1
Cluster 2
Cluster 3
Cluster 4
T exhausted
T prec. exhausted
SLEC
Effectors interm.
Effectors cycling
Early effectors
Memory T precursors
E
Natural Killer cell signature
Score
Module Score
Cluster
Early effector T cells
Early effector T cells
Memory T precursors
T prec. exhausted cells
KILR T cells
F
Gated on KLRK1+OT-I T cells
Norm. to mode
CD103 (PerCP-Cy5.5)
CD103+ KILR
0.0021
% of KILR cells
DEREG- Effectors
DEREG+ Effectors
DEREG- KILR gate
DEREG+ KILR gate
G
FACS gating
AE
DP
KILR
CD49d
KLRK1
FACS gates illustrated on scRNAdata
Itga4
Klrk1
Gzma
0.026
Gzmb
0.017
Fold change, norm. to GAPDH
Gzmk
0.026
Il2ra
0.026
Il7r
0.009
Cd7
0.016
Ifitm1
0.003
Klrd1
0.012
Cxcr6
0.025
DEREG+
DEREG-
DP CD49d+KLRK1+
KILR CD49d-KLRK1+
AE CD49d+KLRK1-

**Figure 5 – Figure supplement 1**

**A-E.** Additional analysis for the scRNAseq experiment shown in Figure 5A-H. **A.** Gene set enrichment analyses comparing the ranked expression of IL-2 induced genes in DEREG$^{+}$ versus DEREG$^{-}$ OT-I T-cell samples. The list of IL-2-responsive genes is based on the indicated publications [40, 41]. P-value (shown in italics) is estimated based on an adaptive multi-level split Monte-Carlo scheme. **B.** Violin plot indicating the expression of *Gzmb* in OT-I T cells primed in DEREG$^{-}$ and DEREG$^{+}$ mice. **C.** The intensity of the blue color indicates the level of expression of indicated genes in individual cells. **D.** Projection of clusters of primed OT-I T cells (Figure 5B) on the UMAP visualization of CD8$^{+}$ T cells subsets formed during an anti-viral response using TILPRED algorithm. **E.** NK cell signature was calculated for each cluster. Top: The intensity of the blue color indicates the level of expression of NK signature genes. Bottom: violin plot showing NK cell signature score calculated for each cluster.

**F.** Additional results of the experiments described in Figure 5I. CD103 expression on KLRK1$^{+}$ CD49d$^{+}$ IL-7R$^{-}$ T cells (Effector) and KLRK1$^{+}$ CD49d$^{-}$ IL-7R$^{+}$ T cells (KILR). Left: A representative histogram. Right: percentage of CD103$^{+}$ cells among cells falling into KILR gate. DEREG$^{-}$ n=11, DEREG$^{+}$ n=7. Median is shown. Statistical significance was calculated using two-tailed Mann-Whitney test, p-value is shown in italics.

**G.** Ly5.1 OT-I T cells ($5 \times 10^4$) were transferred into Treg-depleted DEREG$^{+}$ RIP.OVA mice and DEREG$^{-}$ RIP.OVA mice. The next day, mice were immunized with DC-OVA. On day 3 post immunization, three populations of splenic OT-I T cells (identified as Ly5.1$^{+}$ CD8$^{+}$) were FACS-sorted as indicated: CD49d$^{+}$ KLRK1$^{-}$ antigen-experienced (AE), CD49d$^{+}$ KLRK1$^{+}$ double-positive (DP), and CD49d$^{-}$ KLRK1$^{+}$ (KILR). The correspondence of the FACS gates to scRNAseq clusters is shown. Sorted cells were analyzed via RT-qPCR. The expression of indicated genes was normalized to *Gapdh*. Mean + SEM is indicated. Four independent experiments. Statistical significance was calculated by Kruskal-Wallis test, p-value is shown in italics.

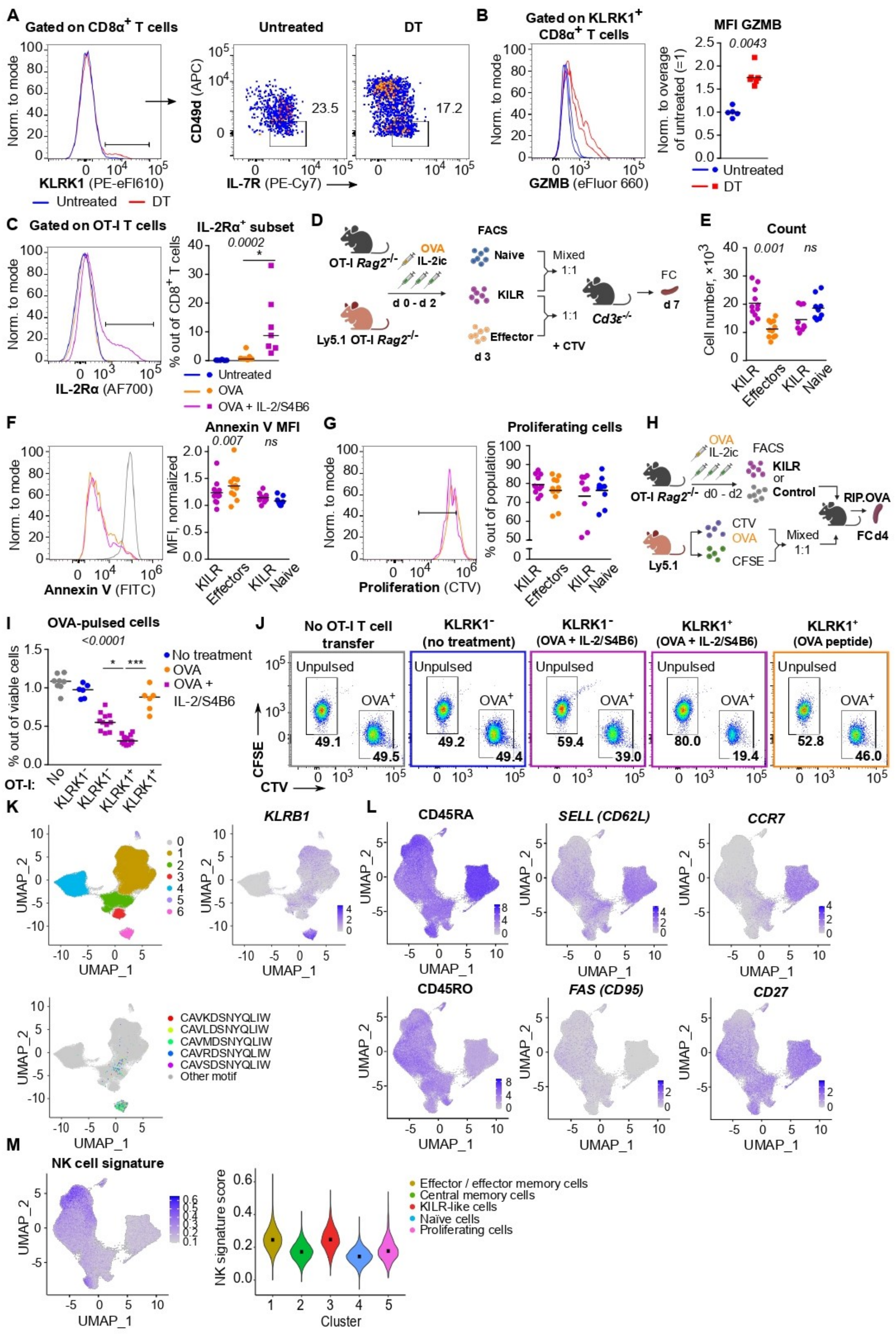
A
Gated on CD8α+ T cells
Norm. to mode
KLRK1 (PE-eFl610)
Untreated
DT
CD49d (APC)
IL-7R (PE-Cy7)
23.5
17.2
B
Gated on KLRK1+ CD8α+ T cells
GZMB (eFluor 660)
MFI GZMB
0.0043
Norm. to overage of untreated (=1)
Untreated
DT
C
Gated on OT-I T cells
IL-2Rα (AF700)
IL-2Rα+ subset
0.0002
% out of CD8+ T cells
Untreated
OVA
OVA + IL-2/S4B6
D
OT-I Rag2-/-
OVA
IL-2ic
d 0 - d 2
Ly5.1 OT-I Rag2-/-
FACS
Naive
KILR
Effector
d 3
Mixed
1:1
+ CTV
Cd3ε-/-
FC
d 7
E
Count
0.001
ns
Cell number, ×10^3
KILR
Effectors
Naive
F
Annexin V (FITC)
Annexin V MFI
0.007
ns
MFI, normalized
G
Proliferation (CTV)
Proliferating cells
% out of population
H
OVA
IL-2ic
FACS
KILR
or
Control
OT-I Rag2-/-
d0 - d2
RIP.OVA
FC d4
Ly5.1
CTV
CFSE
Mixed
1:1
I
OVA-pulsed cells
<0.0001
% out of viable cells
No treatment
OVA
OVA + IL-2/S4B6
OT-I:
No
KLRK1-
KLRK1+
J
No OT-I T cell transfer
KLRK1- (no treatment)
KLRK1- (OVA + IL-2/S4B6)
KLRK1+ (OVA + IL-2/S4B6)
KLRK1+ (OVA peptide)
Unpulsed
OVA+
49.1
49.5
49.2
49.4
59.4
39.0
80.0
19.4
52.8
46.0
CFSE
CTV
K
KLRB1
UMAP_1
UMAP_2
CAVKDSNYQLIW
CAVLDSNYQLIW
CAVMDSNYQLIW
CAVRDSNYQLIW
CAVSDSNYQLIW
Other motif
L
CD45RA
SELL (CD62L)
CCR7
CD45RO
FAS (CD95)
CD27
M
NK cell signature
NK signature score
Cluster
Effector / effector memory cells
Central memory cells
KILR-like cells
Naïve cells
Proliferating cells

**Figure 6 – Figure supplement 1**

**A.** Representative experiment out of three in total for the data shown in Figure 6A.
**B.** GZMB levels in $CD8^+$ T cells from the experiment shown in Figure 6A. Left: A representative histogram. Right: Geometric mean fluorescence intensity (MFI) of anti-GZMB-eFluor 660 antibody on $CD8^+$ T cells was determined. Obtained values were normalized to the average of MFI of untreated samples in each experiment (=1). DT treated n=6, Untreated n=5 mice. Statistical significance was calculated by two-tailed Mann-Whitney test (p-value is shown in italics). Median is shown.
**C**. Additional results of the experiments described in Figure 6B-E. IL-2Rα expression on OT-I T cells from mice treated with OVA, OVA + IL-2/S4B6, and untreated control. Statistical significance was calculated by Kruskal-Wallis test (p-value is shown in italics) with Dunn's multiple comparison post-test (* <0.05). Median is shown.

**D**-**G**. Additional data for the experiment shown in Figure 6F-G. **D**. Scheme of the experiment. KILR T cells were induced in OT-I *Rag2*$^{-/-}$ and congenic Ly5.1 OT-I *Rag2*$^{-/-}$ mice via IL-2/S4B6 and OVA. Three populations of $CD8^+$ OT-I T cells were FACS sorted as: naive ($KLRK1^-$ $CD49d^-$), KILR ($KLRK1^+$ $CD49d^-$ ), effector ($KLRK1^-$ $CD49d^+$). KILR cells were mixed with one of the control populations in 1:1 ratio, stained with CTV proliferation dye, and injected to *CD3ε*$^{-/-}$ recipient mouse. Each mouse got $400 \times 10^3$, or $500 \times 10^3$ cells in total in two independent experiments (KILR + Effector n=11, KILR + Naïve n=9). On day 7 of the experiment (three days post transfer) spleens of the recipients were analyzed by flow cytometry. **E**. Number of cells originating from transferred KILR, effector, and naïve cells, per spleens of recipient. Mean is shown. **F**. Annexin V staining, representative histogram (positive control in grey) on left, normalized geometric mean fluorescent intensity (MFI) on right. Statistical significance was calculated by Wilcoxon matched-pairs signed rank test (p-value is shown in italics), ns – not significant (>0.05). Mean is shown. **G**. Proliferating cells, gated as $CTV^{low}$ cells (2 and more divisions). Left: representative staining. Right: % of proliferating cells out of total cell population. Mean is shown.
**H-J.** Additional data for the experiment shown in Figure 6H. **H.** Scheme of the experiment. OT-I *Rag2*$^{-/-}$ mice were treated with OVA peptide (day 0), and/or IL-2/S4B6 (days 0, 1 and 2) or left untreated. On day 3, $KLRK1^+$ $CD8^+$ or $KLRK1^-$ $CD8^+$ OT-I T cells were FACS sorted and transferred to recipient RIP.OVA mice, which have received a mixture of OVA-pulsed CTV-loaded and unpulsed CFSE-loaded target cells obtained from Ly5.1 mice at ~1:1 ratio earlier the same day. Spleens were analyzed by flow cytometry on day 4 of the experiment. **I.** Percentage of OVA-pulsed target cells out of all viable splenocytes. No OT-I T cell transfer n=8, $KLRK1^-$ (no treatment) n=6, $KLRK1^-$ (OVA + IL-2/JES6) n=11, $KLRK1^+$ (OVA + IL-2/JES6) n=13, $KLRK1^+$ (OVA peptide) n=6. **J.** A representative experiment out of 4 in total. Statistical significance was calculated by Kruskal-Wallis test (p-value is shown in italics) with Dunn's multiple comparison post-test (* <0.05, ***<0.001). Median is shown.
**K.** Identification of MAIT cells in the Human $CD8^+$ T cell atlas. Top left: UMAP projection. Individual cells are assigned to individual clusters based on unsupervised clustering. Cluster 6 corresponds to MAIT cells. Top right: The intensity of the blue color indicates the level of *KLRB1* expression (a MAIT cell marker) in individual cells. Bottom: Cells expressing five most abundant semi-invariant MAIT cell-specific receptor-alpha (TCRα) are shown.
**L.** The intensity of the blue color indicates the expression level of selected genes based on the detection using hash-tagged antibodies (CD45RA and CD45RO) or transcripts in Human $CD8^+$ T cell atlas after the removal of MAIT cells.

**M.** NK cell signature was calculated for each cluster of the Human $CD8^+$ T cell atlas. Left: The intensity of the blue color indicates the level of expression of NK signature genes. Right: violin plot showing NK cell signature score calculated for each cluster.

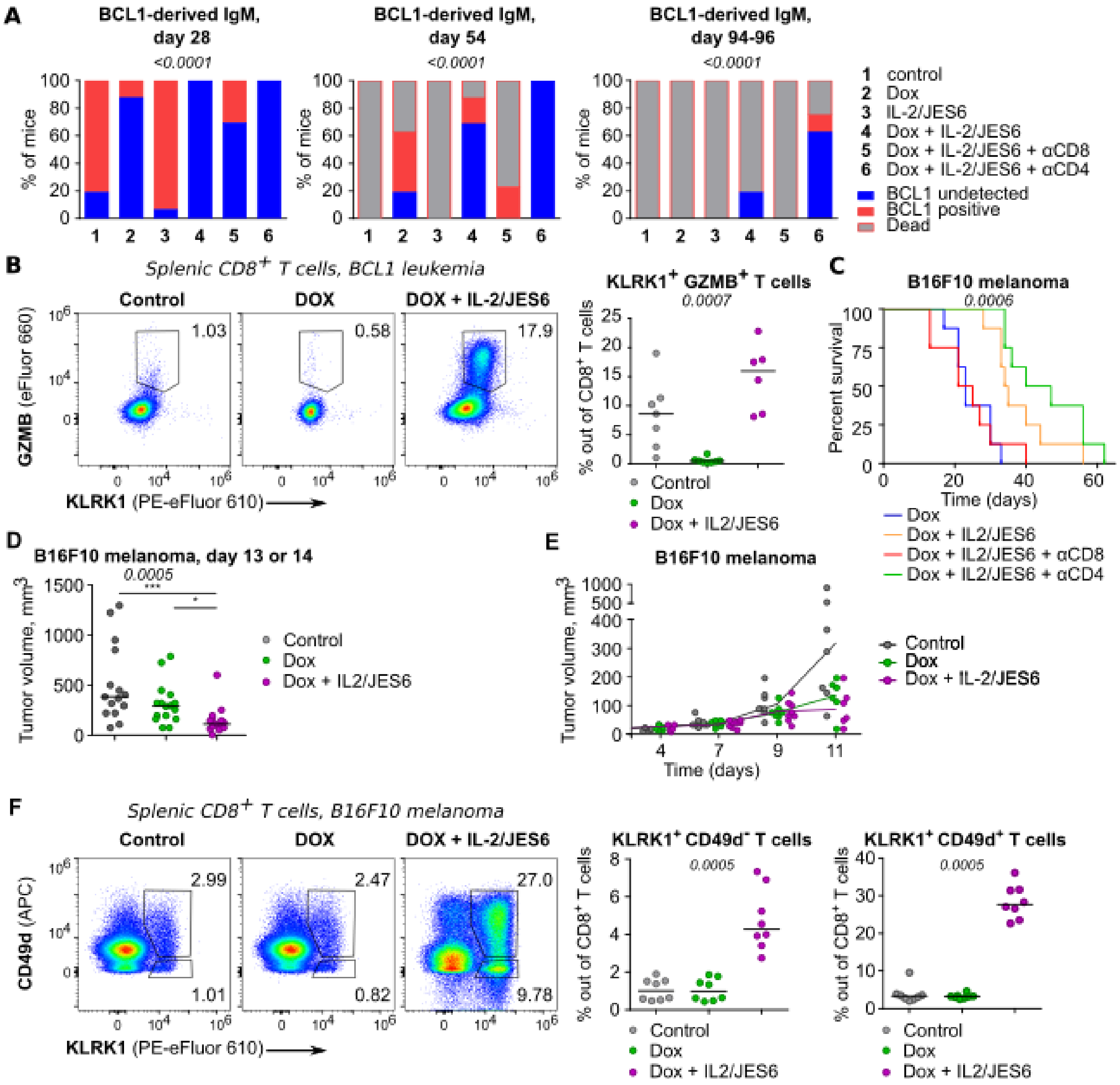


**Figure 7 – Figure supplement 1**

**A.** In experiments described in Figure 7A-B, BCL1 IgM/IgD in the serum on days 28, 54, and 94-96 post tumor inoculation was detected by ELISA. n=16 mice per group in two independent experiments with two exceptions: (i) "Dox + IL-2/JES6 + αCD4" conditions on day 54 where the analysis failed for the second experiment and thus, n=8 mice and (ii) control conditions on day 54, when one mouse was not analyzed for technical reasons and thus, n=15. The statistical significance was calculated using Chi-square test (two groups of the outcomes: mice with undetectable BCL1 vs. mice with detectable BCL1 or dead), p-value is shown in italics.

**B.** Addition data for the experiment shown in Figure 7C. Expression of KLRK1 and GZMB on splenic $CD8^+$ T cells. Left: representative staining. Right: percentage of $KLRK1^+$ $GZMB^+$ cells out of total $CD8^+$ T cells. Kruskal-Wallis test, p-value is shown in italics, median is shown.

**C**. On day 0, C57Bl/6J mice were inoculated with B16F10 melanoma cells. On day 6, mice received doxorubicin (Dox) with/without anti-CD4 or anti-CD8 depletion mAb. On three consecutive days, mice were treated with IL-2/JES6 (see scheme Figure 7D). Survival curves. n=8 mice per group in one experiments. The data shown here partially overlap with data shown in Figure 7E. There were two experiments in total, but only the second experiment had the mouse groups with the depletion of $CD8^+$ or $CD4^+$ cells. Figure 7E shows pooled data for all conditions included in both experiments. Here, results from the second experiment are shown. Statistical significance was calculated by Log-rank (Mantel-Cox) test (p-value is shown in italics).

**D**. Tumor volume recorded on day 13 or 14 in the experiments shown in Figure 7E. Kruskal-Wallis test (p-value is shown in italics) with Dunn's multiple comparison post-test (* <0.05, ***<0.001), median is shown. Control n=15, Dox n=16, Dox + IL-2/JES6 n=15 in two independent experiments.

**E**. Tumor volume growth in the experiment shown in Figure 7F-H. On day 11, tumors were harvested for flow cytometry analysis. n=8 per group in two independent experiments.

**F**. Addition data for the experiment shown in Figure 7F-H. Expression of KLRK1 and CD49d on splenic $CD8^+$ T cells. Left: representative staining. Right: percentage of $KLRK1^+$ $CD49d^-$ and $KLRK1^+$ $CD49d^+$ cells out of total $CD8^+$ T cells. Kruskal-Wallis test (p-value is shown in italics), median is shown.

**Supplementary file Table S1**

Differentially expressed genes for all clusters shown in Figure 5B. The used differential expression criteria were fold change above 2, minimum of 0.1 difference in the fraction of detection between each cluster and the rest of the cells and adjusted p-value below 0.01. For each gene, the fractions of cells with at least one detected transcript in the tested cluster (pct.1) or among all other cells (pct.2) are listed. The columns 'p_val', 'avg_log2FC' and 'p_val_adj' show p-values (Mann Whitney U Test), log2 fold changes and adjusted p-values (Bonferroni correction).

**Supplementary file Table S2**

Differentially expressed genes between the super-effector-like cell cluster and all the remaining cells in the human $CD8^+$ atlas (after MAIT removal). The used differential expression criteria were average fold change above 1.5 and adjusted p-value below 0.01. For each gene, the fractions of super-effector-like cells (pct.1) or remaining cells (pct.2) with at least one detected transcript are listed. The columns 'p_val ', 'avg_log2FC' and 'p_val_adj' show p-values (Mann Whitney U Test), log2 fold changes and adjusted p-values (Bonferroni correction).
